# Genomic inversion drives small colony variant formation and increased virulence in *P. aeruginosa*

**DOI:** 10.1101/356386

**Authors:** Sharon Irvine, Boyke Bunk, Hannah K. Bayes, Cathrin Sprӧer, James P. R. Connolly, Anne Six, Thomas J. Evans, Andrew J. Roe, Jӧrg Overmann, Daniel Walker

## Abstract

Phenotypic change is a hallmark of bacterial adaptation during chronic infection. In the case of chronic *Pseudomonas aeruginosa* lung infection in patients with cystic fibrosis, well-characterised phenotypic variants include mucoid and small colony variants (SCVs). It has previously been shown that SCVs can be reproducibly isolated from the murine lung following the establishment of chronic infection with mucoid *P. aeruginosa* strain NH57388A. Here we show, using a combination of singlemolecule real-time (PacBio) and Illumina sequencing that the genetic switch for conversion to the SCV phenotype is a large genomic inversion through recombination between homologous regions of two rRNA operons. This phenotypic conversion is associated with large-scale transcriptional changes distributed throughout the genome. This global rewiring of the cellular transcriptomic output results in changes to normally differentially regulated genes that modulate resistance to oxidative stress, central metabolism and virulence. These changes are of clinical relevance since the appearance of SCVs during chronic infection is associated with declining lung function.

## Introduction

Phenotypic variation is a hallmark of adaptation to the host during chronic bacterial infection. There is considerable interest in slow growing subpopulations of bacteria, termed small colony variants (SCVs), due to their association with persistent infections^12^. The SCV variant is common to diverse bacteria and is characterised by phenotypes including reduced growth, increased biofilm production^3^, antibiotic resistance and hyperpiliation. SCVs have been described for a wide range of bacterial genera and species including *Staphylococcus aureus*^4,5^, *Staphylococcus epidermidis*^6^, *Streptococcus* sp^7,8^, *Enterococcus*^9^, *Listeria*^10^, *Burkholderia*^11^, *Salmonella*^12^, *Vibrio*^13^, *Shigella, Brucella*^14^, *Lactobacillus, Serratia and Neisseria*^15^. In the case of *Pseudomonas aeruginosa*, SCVs are commonly associated with chronic infection of the lung in patients with cystic fibrosis^16-21^.

*P. aeruginosa* is the major proven cause of mortality in patients with CF and chronic infection leads to a progressive decline in pulmonary function and inevitably respiratory failure^22^. Despite intensive anti-pseudomonal chemotherapy greatly improving the prognosis for CF patients^23^, the current median age at death for CF patients is around thirty years in developed countries^24^. The frequent failure of antibiotic therapy and host defences to eradicate *P. aeruginosa* from the CF lung is thought to be largely due to the increased antibiotic tolerance when growing in the biofilm state and the appearance of mucoid phenotyic variants that are a hallmark of adaptation in the chronically infected lung. A further complicating factor is the appearance of highly adherent SCVs that are adept at biofilm formation^18,19,25^. *P. aeruginosa* SCVs may display high intracellular c-di-GMP levels^19,20,26-28^, enhanced biofilm formation, high fimbrial expression, repression of flagellar genes, resistance to phagocytosis and enhanced antibiotic resistance. Most importantly, the appearance of SCVs in the CF lung correlates with poor patient clinical outcome^11,17,29-31^.

There are a range of genetic changes that have been shown to be responsible for the phenotypic switch to the SCV phenotype in *P. aeruginosa*, including mutations in the Wsp system and *yfiBNR* operon that form part of the c-di-GMP regulatory system in *P.* aeruginosa^32-36^. However, identification of the major clinically relevant pathways of conversion to the SCV phenotype are complicated by the unstable phenotype displayed by many SCVs with reversion to a normal colony phenotype frequently observed preventing successful comparative genetic studies on clinical SCVs and their closely related parent strains. In the case of S. *aureus*, which also forms clinically relevant SCVs, recent work has shown that a reversible large scale chromosomal inversion is the genetic basis of the switch between a normal colony and SCV isolated from the same patient^37^. In addition, *S. aureus* SCVs, which are commonly isolated from the CF lung, are highly resistant to oxidative stress, suggesting that conversion to the SCV phenotype may be an adaptation to environment present in chronically inflamed host tissue^38^.

In this work, we have determined the genetic basis of phenotypic conversion from the mucoid to the SCV phenotype for SCVs isolated from the chronic lung infection model described by Bayes et al 2016^39^. For two SCVs isolated from this work, we have shown through a combination of single-molecule real-time (SMRT, Pacific Biosciences) and Illumina sequencing that a large and stable chromosomal inversion accompanies conversion to the SCV phenotype. Genome inversion is accompanied by transcriptional changes to a large number of genes that most notably includes downregulation of a number of genes encoding metabolic enzymes, DNA repair proteins and heat shock proteins and the upregulation of genes encoding proteins involved in the response to oxidative stress. The absence of other obvious genetic change suggests that this chromosomal inversion is the genetic basis of conversion to the SCV phenotype.

## Results

*P. aeruginosa* SCVs are commonly isolated from patients with cystic fibrosis and have been isolated *in vitro* as well as from experimental infection models following aminoglycoside treatment^18,31,40^. The work of Bayes et al describes the isolation and partial characterisation of SCVs isolated from a chronic murine *P. aeruginosa* lung infection model^39^. In this model, animals were inoculated with *P. aeruginosa* strain NH57388A (NHMuc) a mucoid clinical isolate, embedded in agar beads. NHMuc has a known mutation in the gene encoding the anti-sigma factor MucA, that results in alginate overproduction^41,42^. Recovered bacteria from lung homogenate samples display two distinct colony morphologies: typical large mucoid colonies identical in morphology to the inoculating strain and SCVs. Mucoid colonies were evident after 24 hours of growth on agar plates at 37°C with SCVs visible only after 48 hours of growth on agar plates^39^.

To understand the genetic basis of this phenotypic change, we initially performed Illumina HiSeq whole-genome sequencing and genomic comparison between NH and two separate SCVs (SCVJan and SCVFeb) isolated from independent in vivo experiments. However, despite their gross phenotypic differences, this analysis failed to identify any genetic differences between the SCVs and the parent strain.

Next, we utilised the ultra-long reads produced by single-molecule real-time PacBio sequencing to attempt to identify any large scale genome rearrangements that could drive conversion to the SCV phenotype. Using this technique we identified a large scale genomic inversion accompanying conversion from the parent mucoid to SCV phenotype in both SCVJan and SCVFeb. Closer inspection of the genome sequence identified the start and end points of the inversion which for both SCVJan and SCVFeb begins at the first rRNA operon (0.72 Mbp) and ends at the third rRNA operon (5.21 Mbp). Exact chromosomal breakpoints were identified in the corresponding 16S rRNA genes by performing a MAUVE breakpoint analysis (Figure 1). Furthermore, genome analysis revealed a 250 bp shortened 16S rRNA gene (16St) in both SCV strains, which is reflected in the reduced genome sizes of the SCVs (SCVJan 6,213,026, SCVFeb 6,213,029; Figure 2a) compared to the parent strain, NHmuc (6,213,276 bp; Figure 2b). There were no further differences in the number of protein coding genes (5619), rRNAs (12) or tRNAs (57) between SCVs and the parent strain. No SNPs could be identified in protein coding genes. A similar genome inversion was not identified (using a PCR based strategy) in a mucoid strain (NHMucJan) that was phenotypically identical to the parent strain (NHMuc) and isolated from the same chronic infection model as SCVJan (Figure S1). Interestingly, comparison of the SCVJan and SCVFeb genomes with that of an SCV (SCV20265) isolated from a CF patient^43^, which has recently been sequenced by PacBio sequencing, revealed an almost identical chromosomal inversion. However, in the case of SCV20265 the inversion was not accompanied by truncation of the 16S rRNA gene in the third rRNA operon (Figure 1).

**Figure 1.**
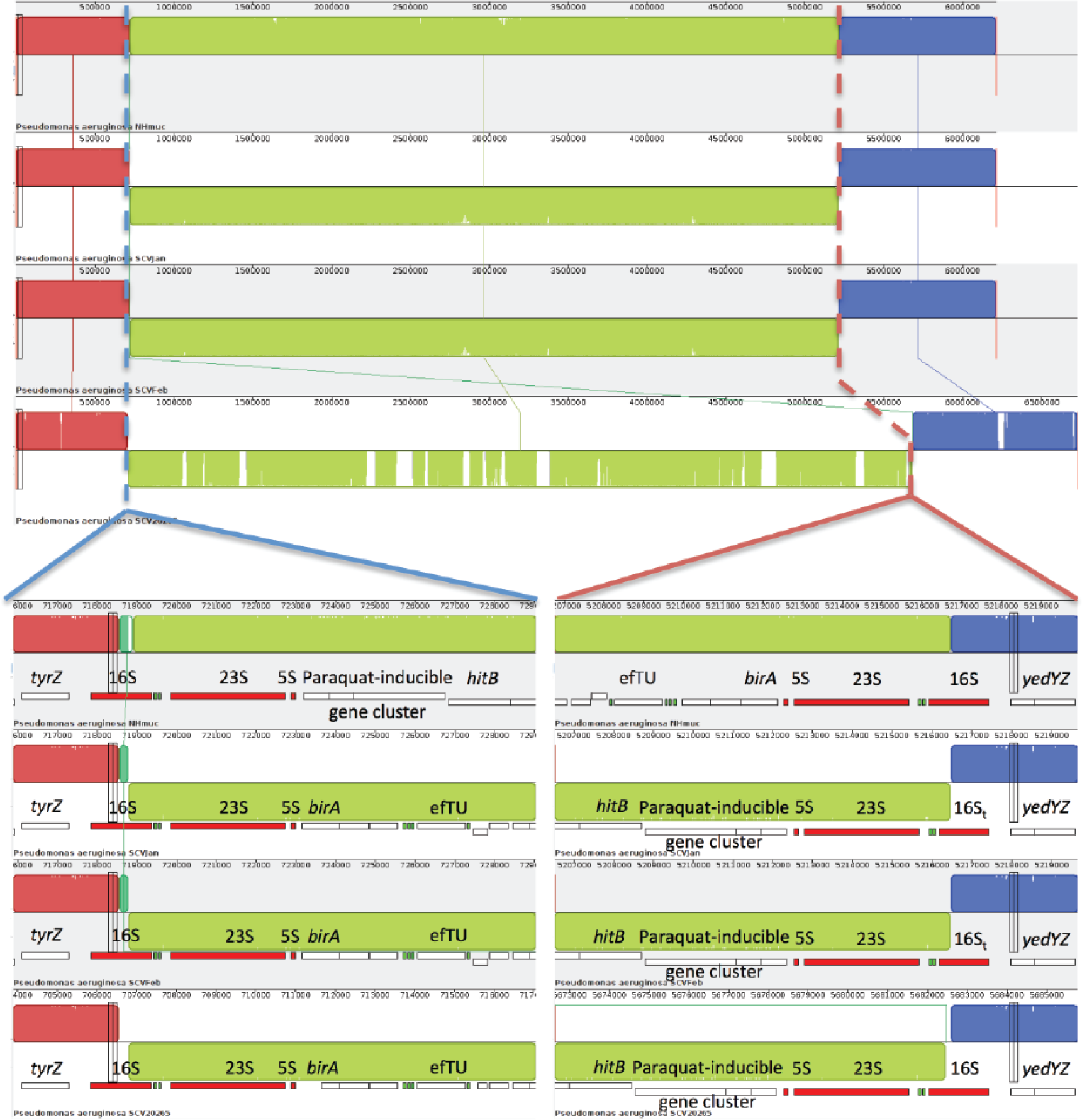
A common large scale chromosomal inversion in three *P. aeruginosa* strains is the genetic basis of conversion to the SCV phenotype. From top to bottom strains NHMuC, SCVJan, SCVFeb and SCV20265 are displayed. Dashed lines indicate the inversion breakpoints present in the 16S rRNA genes. An inversion with highly similar breakpoints is present in the genome of strain SCV20265 a SCV isolated from a patient with CF. Within strains SCVJan and SCVFeb a unique truncated version of the 16S rRNA gene (16St) could be resolved, which could not be detected in strain SCV20265.

**Figure 2.**
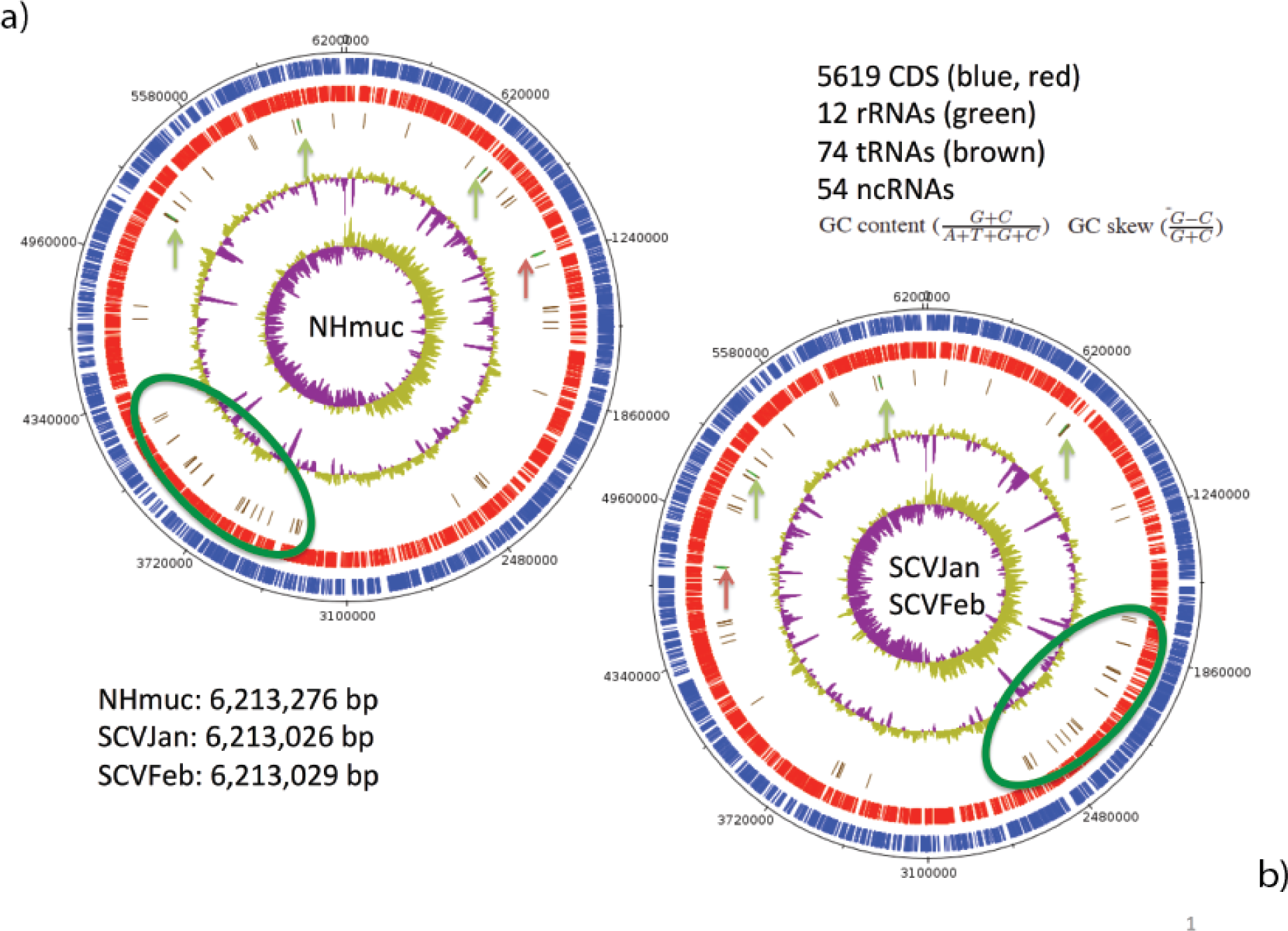
Chromosomal maps of *P. aeruginosa* NHmuc (a) and SCVJan/SCVFeb (b). The circular genomes of both strains are shown. Genomes of both SCV strains are 250 bp smaller compared to the parental strain NHmuc. Exact genome sizes are given in lower left corner. In blue (circle1) genes lying on the forward strand are shown and in red (circle 2) those on the reverse strand. In circle 3 tRNA genes are shown in brown, often clustered together with green rRNA genes, which have been additionally marked by vertical arrows. The red arrow shows the transposition of rRNA operon 3 in addition to that of a large tRNA region (green ellipse) due to the described chromosomal inversion. Circle 4 shows the GC content, whereas in circle 5 a GC skew is shown. Number of CDS, rRNAs, tRNAs and ncRNAs are identical in all strains (upper right corner according to GenBank submission). This map has been created using DNAplotter^72^ (Carver *et al.*, 2009).

## Transcriptional and phenotypic changes on conversion to the SCV phenotype

To determine the transcriptional changes associated with conversion to the SCV phenotype, we performed RNA sequencing (RNA-Seq) analysis of the parent strain, NHMuc, and two SCV strains grown in LB broth. Initial analysis showed that SCVJan and SCVFeb have highly similar gene expression profiles that are distinct from that of NHMuc. RNA-Seq data for all strains was collected in triplicate and data for SCVJan and SCVFeb were combined to compare with NHMuc. Relative to NHMuc, 190 genes showed >2-fold upregulation and 364 genes showed >2-fold downregulation in SCVJan/SCVFeb (Table S1). Interestingly, the transcriptional changes associated with genomic inversion and that drive conversion to the SCV phenotype are not restricted to genes close to or within the inversion breakpoints, with major upregulated and downregulated genes distributed relatively evenly throughout the genome (Figure 3).

**Figure 3.**
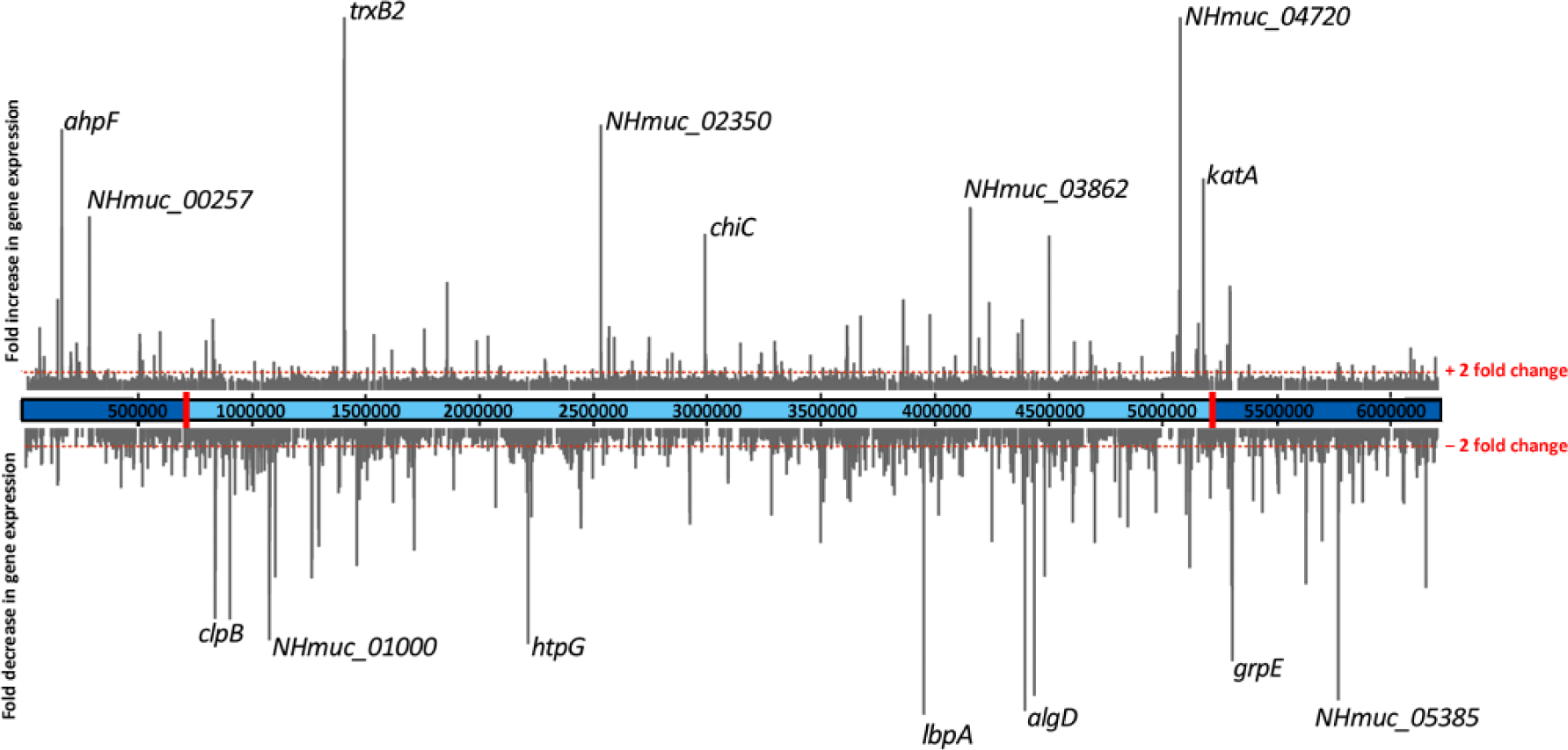
Global changes in gene transcription on conversion to the SCV phenotype. Fold-changes in gene expression for SCVJan/Feb relative to NHMuc are shown in the context of the NHMuc genome. Breakpoints that define the genomic inversion present in SCVJan and SCVFeb are indicated by red rectangles.

Major functional classes of genes downregulated in SCVJan/SCVFeb include those involved in energy metabolism, amino acid and protein biosynthesis, DNA replication and recombination and cell wall/LPS/capsule biosynthesis, which together are consistent with the slow growth rate observed for SCVs. Notably, genes encoding heat shock proteins and other molecular chaperones (IbpA, GrpE, HtpG, ClpB, DnaK, GroES, DnaJ and ClpX) are highly represented among the most strongly downregulated genes in the SCVs (Table 1). Conversely, genes that function in the response to oxidative stress and those that encode secreted virulence factors are largely upregulated in SCVJan/SCVFeb. Indeed, five of the ten most highly upregulated genes in SCVJan/SCVFeb are those associated with the response to oxidative stress (Table 2). Highly upregulated oxidative stress genes include, *katA^44^* which encodes the major catalase of *P. aeruginosa, ahpB, ahpC* and ahpF^45-47^ that encode subunits of alkyl hydroperoxide reductase and *trxB2* that encodes thioredoxin reductase 2^48-50^. Consistent with the observed transcriptional changes, catalase activity was strongly increased in SCVJan relative to NHMuc (Figure 4A).

**Figure 4.**
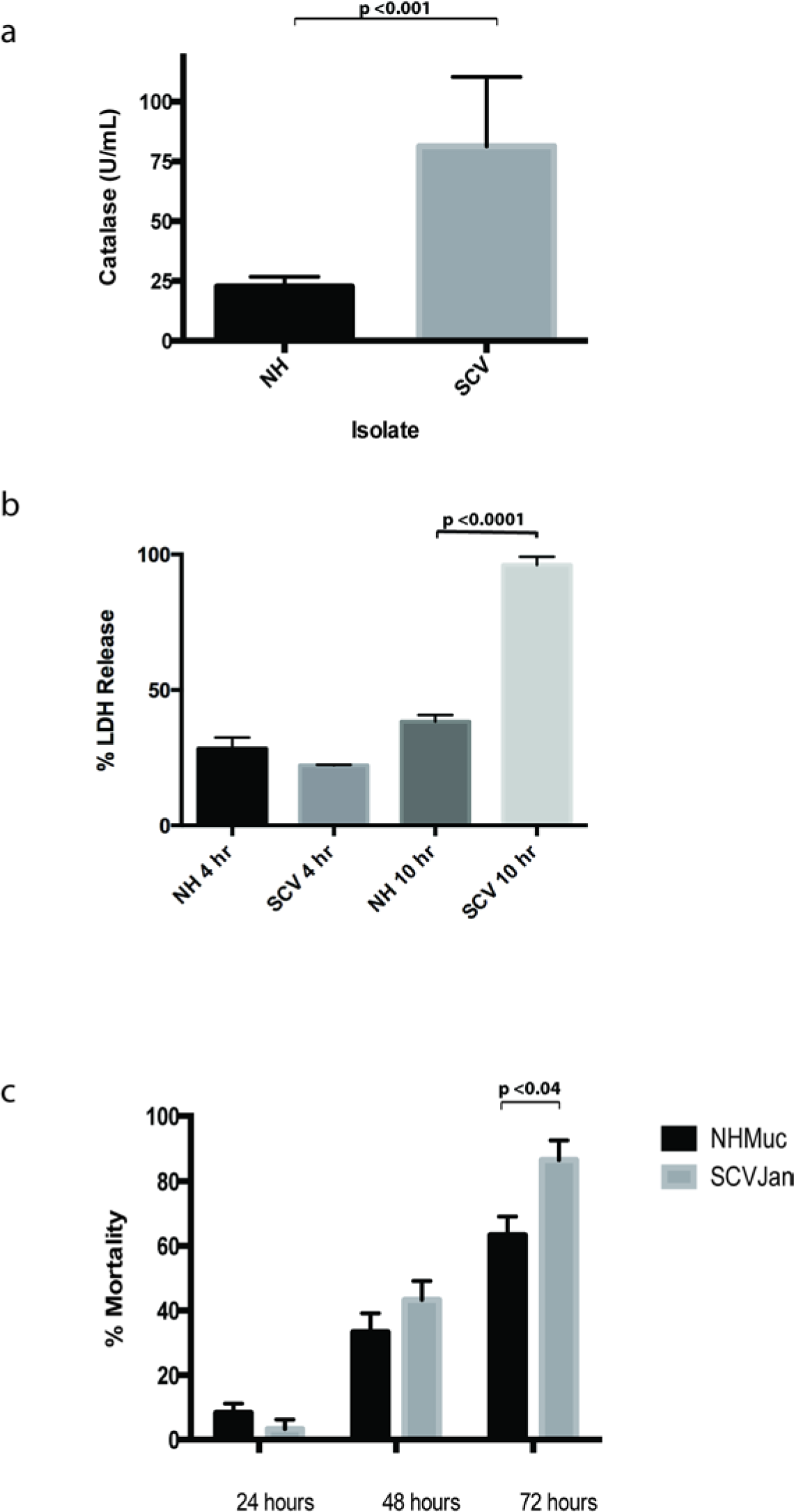
Phenotypic characterisation of SCVJan. a) Catalase activity assay demonstrating marked increase in catalase activity in the SCV as compared to the NH parent strain. b) % LDH released from macrophage cell line with comparison between NHMuc and SCVJan over a 4 and 10 hour time period. c) *Galleria mellonella* larvae survival over time when infected with the NHMuc and SCVJan strains monitored over 72 hours.

**Table 1:** Highly downregulated genes in SCVs relative to NHMuc

| Gene ID | Fold Change<sup>a</sup> | p-Value<sup>b</sup> | Protein description | PAO1 gene ID |
| --- | --- | --- | --- | --- |
| NHMuc_01025 | -75 | $2.6 \times 10^{-11}$ | Putative acetyltransferase | PA4364 |
| mexC | -53 | $1.1 \times 10^{-11}$ | RND multidrug efflux fusion protein MexC | PA4599 |
| ibpA | -31 | $1.1 \times 10^{-7}$ | Heat-shock protein IbpA | PA3126 |
| algD | -30 | $1.0 \times 10^{-9}$ | GDP-mannose 6-dehydrogenase AlgD | PA3540 |
| NHMuc_05385 | -29 | $8.5 \times 10^{-8}$ | 17 kDa surface antigen | PA5182 |
| NHMuc_04138 | -29 | $1.3 \times 10^{-10}$ | Periplasmic metal-binding protein | PA3520 |
| grpE | -25 | $1.2 \times 10^{-7}$ | Heat shock protein GrpE | PA4762 |
| htpG | -23 | $3.6 \times 10^{-7}$ | Heat shock protein 90 | PA1596 |
| NHMuc_0100 | -23 | $1.8 \times 10^{-5}$ | Hypothetical, unclassified, unknown | |
| clpB | -21 | $6.5 \times 10^{-6}$ | ClpB protein | PA4542 |
| nfxB | -21 | $3.7 \times 10^{-7}$ | Transcriptional regulator NfxB | PA4600 |
| fsxA | -18 | $7.9 \times 10^{-8}$ | FxsA protein | PA4387 |
| NHMuc_04950 | -18 | $3.3 \times 10^{-6}$ | Molecular chaperone DnaK | PA4761 |
| NHMuc_05744 | -17 | $5.8 \times 10^{-5}$ | Putative lipoprotein | PA5526 |
| dapB | -17 | $2.8 \times 10^{-6}$ | Dihydrodipicolinate reductase | PA4759 |
| hslV | -17 | $4.1 \times 10^{-7}$ | ATP-dependent protease subunit | PA5053 |
| NHMuc_01173 | -16 | $4.7 \times 10^{-6}$ | Hypothetical protein | PA2756 |
| NHMuc_01024 | -16 | $6.1 \times 10^{-7}$ | Putative transporter | PA4365 |
| NHMuc_04180 | -16 | $3.3 \times 10^{-6}$ | Recombinase A | PA3617 |
| mexD | -15 | $9.5 \times 10^{-7}$ | RND multidrug efflux transporter MexD | PA4598 |
| NHMuc_04776 | -15 | $6.1 \times 10^{-6}$ | PAS/PAC sensor signal transduction histidine kinase | PA4197 |
| rsmA | -15 | $7.5 \times 10^{-4}$ | Carbon storage regulator | PA0905 |
| NHMuc_01174 | -14 | $3.3 \times 10^{-7}$ | Hypothetical protein | PA0737 |
| NHMuc_01594 | -13 | $2.1 \times 10^{-5}$ | Putative oxidoreductase | PA1137 |
| mucA | -13 | $7.4 \times 10^{-5}$ | Anti-sigma factor MucA | PA0763 |
| NHMuc_03262 | -12 | $2.5 \times 10^{-9}$ | Hypothetical protein | PA3505 |
| NHMuc_04386 | -12 | $6.8 \times 10^{-5}$ | Surface antigen | PA3819 |
| amrZ | -12 | $3.0 \times 10^{-5}$ | Alginate and motility regulator Z | PA3385 |
| glnA | -12 | $5.8 \times 10^{-6}$ | Gutamine synthetase | PA5119 |
| groES | -12 | $3.9 \times 10^{-5}$ | Co-chaperonin GroES | PA4386 |
| dnaJ | -12 | $7.5 \times 10^{-5}$ | Chaperone protein DnaJ | PA4760 |
| algU | -11 | $1.8 \times 10^{-4}$ | RNA polymerase sigma factor AlgU | PA0762 |
| clpX | -11 | $4.5 \times 10^{-4}$ | ATP-dependent protease subunit ClpX | PA1802 |
| NHMuc_04521 | -11 | $1.9 \times 10^{-6}$ | Periplasmic ligand-binding sensor | PA3952 |

<sup>a</sup> The magnitude of gene expression (fold change) was determined by comparing transcription in three replicates of NH with that in three replicates each of the two SCV strains.
<sup>b</sup> p-values were assessed by performing an EDGE test using CLC software.

**Table 2:** Highly upregulated genes in SCVs relative to NHMuc

| Gene ID | Fold Change<sup>a</sup> | p-Value<sup>b</sup> | Protein description |
| --- | --- | --- | --- |
| NHMuc_04720 | 62 | $3.2 \times 10^{-16}$ | Hypothetical protein |
| trxB2 | 55 | $4.0 \times 10^{-16}$ | Thioredoxin reductase 2 |
| NHMuc_01300 | 35 | $3.7 \times 10^{-15}$ | Putative alkyl hydroperoxide reductase |
| NHMuc_02350 | 28 | $1.4 \times 10^{-26}$ | Putative acyl carrier protein |
| ahpF | 28 | $8.4 \times 10^{-13}$ | Alkyl hydroperoxide reductase subunit F |
| kata | 23 | $7.1 \times 10^{-10}$ | Catalase |
| NHMuc_03862 | 20 | $6.9 \times 10^{-21}$ | Putative ankyrin domain-containing protein |
| NHMuc_03863 | 19 | $1.8 \times 10^{-18}$ | Putative hydrolase |
| NHMuc_00257 | 19 | $1.4 \times 10^{-12}$ | Putative CBS domain protein |
| ahpC | 18 | $9.0 \times 10^{-8}$ | Alkyl hydroperoxide reductase subunit C |
| chiC | 17 | $1.5 \times 10^{-9}$ | Chitinase |
| NHMuc_04185 | 17 | $6.7 \times 10^{-6}$ | RNA polymerase sigma factor RpoS |
| aprA | 12 | $5.1 \times 10^{-8}$ | Alkaline metalloproteinase |
| NHMuc_04924 | 11 | $3.2 \times 10^{-16}$ | CsbD family protein |
| NHMuc_04718 | 11 | $1.5 \times 10^{-11}$ | Hypothetical protein |
| NHMuc_04925 | 11 | $2.2 \times 10^{-8}$ | Transport-associated |
| NHMuc_00127 | 10 | $5.9 \times 10^{-7}$ | Putative hemolysin |
| Snr1 | 10 | $4.9 \times 10^{-7}$ | Cytochrome c Snr1 |
| lecB | 9 | $5.9 \times 10^{-8}$ | Fucose-binding lectin PA-IIL |
| NHMuc_03413 | 8 | $9.1 \times 10^{-15}$ | Phage terminase, small subunit |
| katB | 8 | $9.1 \times 10^{-6}$ | Catalase |
| NHMuc_04074 | 8 | $2.2 \times 10^{-12}$ | Leucyl-tRNA synthetase |
| phzG2_2 | 7 | $3.3 \times 10^{-6}$ | Pyridoxamine 5'-phosphate oxidase |
| NHMuc_03358 | 7 | $2.0 \times 10^{-8}$ | Putative protein associated inclusion bodies |
| phzE1_1 | 7 | $3.2 \times 10^{-6}$ | Phenazine biosynthesis protein PhzE |
| NHMuc_00055 | 7 | $6.1 \times 10^{-7}$ | Hypothetical protein |
| NHMuc_01628 | 7 | $1.3 \times 10^{-8}$ | Hypothetical protein |
| cbpD | 7 | $4.1 \times 10^{-6}$ | Chitin-binding protein CbpD |
| NHMuc_00546 | 6 | $2.0 \times 10^{-3}$ | LysR transcriptional regulator |
| rhIR | 6 | $1.2 \times 10^{-4}$ | Transcriptional regulator RhIR |
| gcdH | 6 | $4.3 \times 10^{-5}$ | Glutaryl-CoA dehydrogenase |
| NHMuc_01422 | 6 | $4.1 \times 10^{-4}$ | Putative dna-binding stress protein |
| NHMuc_04078 | 6 | $1.5 \times 10^{-10}$ | Oxidoreductase probably involved in sulfite reduction |
| rsaL | 6 | $1.2 \times 10^{-2}$ | Regulatory protein RsaL |

<sup>a</sup> The magnitude of gene expression (fold change) was determined by comparing transcription in three replicates of NH with that in three replicates each of the two SCV strains.
<sup>b</sup> p-values were assessed by performing an EDGE test using CLC software.

Genes encoding a number of secreted virulence factors such as the proteases LasA, LasB and AprA, the frucose-binding lectin LecB^51^ and the chitin binding protein CbpD and chitinase ChiC^52^ were also highly upregulated. Similarly genes encoding hydrogen cyanide synthase and a number of enzymes that function in phenazine biosynthesis are also upregulated (Table S1)^28,53-57^. Phenazines have previously been shown to enhance killing of *Caenorhabditis elegans* by *P. aeruginosa*^58^. The apparent increase in the production of virulence factors by SCVJan/SCVFeb relative to NHMuc suggests increased virulence of the SCV. To directly test this we used an infection model based on infection of the murine macrophage cell line J774A. 1. Cell death of J774A. 1 through LDH release was measured at 4 and 10 hours post infection with NHMuc and SCVJan. At 4 hours, levels of LDH release were similar for NHMuc and SCVJan, whereas at 10 hours LDH release was significantly increased for SCVJan; 96% vs 38%, p < 0.0001 (Figure 4B).

To determine if the increased virulence of the SCV observed against a murine cell line translated to increased virulence in an animal model of infection, we utilised an invertebrate model of infection utilising the larva of the wax moth *Galleria mellonella.* Similar to the macrophage infection assay, SCVJan displayed increased virulence in the *G. mellonella* infection model. Mortality of larvae was measured at 24, 48 and 72 hours post infection. No significant differences in mortality were detected at 24 or 48 hours, whereas at 72 hours % mortality was 86% and 63% (p < 0.04) for SCVJan and NHMuc infected larvae, respectively (Figure 4C). Data from both infection models shows that SCVJan shows increased virulence, relative to NHMuc, which is consistent with the phenotype of SCVs obtained from the human host.^31,59^

## Discussion

In this work we show that *P. aeruginosa* SCVs isolated from a chronic murine lung infection model show a general upregulation of virulence associated genes, relative to the mucoid parent strain, and increased virulence which may begin to explain the link between the appearance of SCVs in chronic lung infection and the associated decline in lung function^60^. In addition, the upregulation of genes that mediate the response to oxidative stress immediately suggests why the isolated SCVs are rapidly selected for in a chronic infection model in which the host immune system is strongly activated.

A key strength of our study was the availability of the parent strain used to establish infection for sequencing. This allowed a meaningful comparative genetic analysis to be performed enabling the determination for the genetic basis of conversion to the SCV phenotype. Surprisingly, SNPs and INDELs were not identified in the SCV genome by Illumina sequencing and single-molecule realtime sequencing was subsequently used to show that the two sequenced SCVs carried a large genomic inversion within 16S rRNA genes. Interestingly, the transcriptional changes associated with genomic inversion are not restricted to genes close to or within the inversion breakpoints, but distributed relatively evenly throughout the genome. Instead the major changes in gene expression are largely restricted to specific functional classes of genes including those that mediate the response to oxidative stress, virulence, DNA repair and recombination, the chaperone network and metabolism. This global rewiring of the cellular transcriptomic output results in concerted transcriptional changes to these normally differentially regulated genes. However, the mechanisms that underlie these transcriptional changes are not clear. A recent study in *E. coli* suggests that position effects on gene expression may be due to local differences in chromosomal structuring and organization, with DNA gyrase playing an important role at certain high activity locations. Further studies will be needed to clarify such positional effects on gene expression in the SCV studied here^61^.

The mechanistic details of how the observed genomic inversion lead to the coordinated expression changes observed here is currently not known, but the observation that a clinical SCV strain (SCV20265) obtained from the lung of a CF patient strain recently sequenced using the same strategy of combining SMRT and Illumina sequencing possesses a similar 16S rRNA based inversion, indicates that this inversion may be a highly clinically relevant route to the SCV phenotype (Figure 1)^43^. Other large scale genome rearrangements including large chromosomal inversions have previously been described in *P. aeruginosa* but these were not associated with conversion to the SCV phenotype^62,63^. A reversible genomic inversion has also recently been shown to mediate the reversible conversion between normal colony and SCV phenotypes in S. aureus^37^. However, in the case of the SCVs isolated in our work the SCV phenotype is stable and revertants to the parent phenotype were not observed. A possible explanation for this observation is that a number of genes encoding proteins involved in DNA repair and recombination, including RecA, are downregulated in the SCV relative to the parent strain (Table 1).

In conclusion, we have shown that a *P. aeruginosa* SCV originated in the lungs of an animal with chronic colonization appears to result from a large chromosomal inversion and associated large-scale transcriptional changes. SCVs have a clear selective advantage in the context of the CF lung, and a better understanding of the drivers that produce the genomic rearrangement observed in this study may provide alternative therapeutic approaches to prevent the appearance of such damaging phenotypic variants.

## Materials and Methods

### Genome assembly and annotation

Purified bacterial genomic DNA was prepared for sequencing on Illumina HiSeq using QIAGEN DNeasy Blood and Tissue Kit as per manufacturers protocol. Sequencing and initial bioinformatics were performed in the Centre for Genomic Research, University of Liverpool. Sequencing reads were mapped to the corresponding reference genome (annotated NH strain). SMRTbell™ template libraries were prepared according to the instructions from Pacific Biosciences, Menlo Park, CA, USA, following the Procedure & Checklist Greater than 10 kb Template Preparation and Sequencing. Briefly, for preparation of 10kb libraries ~10μg genomic DNA isolated from SCVJan, SCVFeb and NHmuc was sheared using g-tubes™ from Covaris, Woburn, MA, USA according to the manufacturer’s instructions. 5-10μg sheared genomic DNA was end-repaired and ligated overnight to hairpin adapters applying components from the DNA/Polymerase Binding Kit P4 from Pacific Biosciences, Menlo Park, CA, USA. Reactions were carried out according to the manufacturer’s instructions. SMRTbell™ template was Exonuclease treated for removal of incompletely formed reaction products. Conditions for annealing of sequencing primers and binding of polymerase to purified SMRTbell™ template were assessed with the Calculator in RS Remote, PacificBiosciences, Menlo Park, CA, USA. SMRT sequencing was carried out on the PacBio RSII (PacificBiosciences, Menlo Park, CA, USA) taking one 180-minutes movie for each SMRT cell. In total 6, 6 and 5 SMRT cells were run respectively. Data from each SMRT Cell was assembled independently using the “RS_HGAP_Assembly.3“ protocol included in SMRTPortal version 2.3.0 using default parameters. Each assembly revealed the fully resolved chromosome in one single contig. Each chromosome was circularized independently, particularly artificial redundancies at the ends of the contigs were removed and all chromosomes were additionally adjusted to *dnaA* as the first gene. Validity of each assembly was checked using the “RS_Bridgemapper.1” protocol. For the purpose of this study it has been confirmed for each of the (repetitive) rRNA operons that enough uniquely mapping long read exist spanning the whole repeat structure. Finally, each genome was error-corrected by a mapping of Illumina reads (paired end reads, 100 bp) onto finished genomes using BWA^64^ with subsequent variant calling using VarScan^65^. A consensus concordance of QV60 could be confirmed for all of the three genomes. Finally, all genomes were annotated using Prokka 1.8^66^. All genome sequences were deposited in NCBI GenBank under Accession Numbers CP013477, CP013478 and CP013479. Illumina short read data has been deposited at EMBL-EBI ENA database under study number PRJEB12456. The shortened 16S rRNA gene for strains SCVJan and SCVFeb was confirmed by PacBio assembly as well as BWA mapping of Illumina reads against the final chromosome showing uniquely mapped reads only at that genome position (data not shown).

### Transcriptome analysis

Ribonucleic acid isolation of the samples was performed in triplicate. Bacterial suspensions were grown to early stationary phase to an OD600 of 1.8 in LB broth at 37 in a shaking incubator. 2 ml of each suspension was pelleted at 12000g for 10 minutes. RNA was extracted from samples using a bead beating/chloroform extraction method as previously described^67^. The samples were digested with DNAse I for 1 hour. Bacterial RNA was enriched using MICROBEnrich (Life Technologies) as per protocol. Ribosomal RNA was depleted using Ribo-Zero Magnetic Gold Kit (Epidemiology) (Epicentre) as per manufacturers protocol. The precipitated sample was resuspended in 20 μΙ of RNAse free water. The concentration of RNA was initially determined using Nanodrop followed by an Agilent Bioanalyser. cDNA was generated by using the methods from the Superscript Double-Stranded cDNA Synthesis Kit (Invitrogen) per manufacturer’s instructions.

Transcriptome analysis was performed using CLC workbench version 7.0 and significantly upregulated and downregulated genes in SCVJan/SCVFeb vs NHMuc were identified using the CLC software package. Transcriptome data was deposited at EMBL-EBI ENA database under study number PRJEB12456.

### PCR

Genomic DNA was extracted from 1.5 ml of bacterial culture using the GenElute Bacterial Genomic DNA Kit (Sigma). Extraction was performed following manufacturer recommendations and DNA was eluted into 100 μL of Elute Solution. PCR detection of the inversion was performed with the KAPA polymerase (KAPA Biosystems, KAPA Long-range HotStart PCR Kit) following the manufacturer’s protocol. Two sets of primers were used: birA-F/yedZ-R and birA-F/tyrZ-R (birA-F: CTCACCGGAGTGGAATC, yedZ-R: T GAGCGCTT ACTGCGT GTT CATCCTGG and tyrZ-R: CCATACCGTGCTTATTAATAAGC) with the genomic DNAs of the mucoid and SCV strains recovered from animals amplifying a fragment of 6262 bp or 6960 bp, respectively.

### Galleria mellonella infection model

Larvae were stored on wood chips at 4 °C. Overnight cultures of bacterial strains were grown in LB broth, diluted 1:100 in the same medium and grown to an OD600 of 0.3 to 0.4 as previously described^68,69^. Cultures were centrifuged and pellets were washed twice and resuspended in 10 mM PBS to an OD600 of 0.1. Serial 10-fold dilutions were made in PBS. Five-microliter aliquots of the serial dilutions were injected using a Hamilton syringe into *G. mellonella* larvae, via the hindmost left proleg as previously described^70^. Ten larvae were injected per dilution for each *Pseudomonas* strain tested. Larvae were incubated in 10-cm plates at 37 °C and the number of dead larvae scored 1 to 4 days after infection. For each strain, data from 3 independent experiments were combined. Larvae were considered dead when they displayed no movement in response to touch. A negative control was used in each experiment to monitor the killing due to physical injury or infection by pathogenic contaminants. Time to death was monitored every 24h post infection. In any instance where more than one control larvae died in any given experiment, the data from infected larvae were not used.

### LDH Release/Cytotoxicity Assay

To investigate the effect of the *P. aeruginosa* strains on macrophages, we infected the J774A.1 cells with NH and SCV. Bacteria were grown for 17 hours to stationary phase in LB broth at 37 °C. Immediately prior to infection, the bacteria were diluted to exponential growth phase with culture medium lacking phenol red and the concentration determined by measuring the optical density at 600nm. Cells were grown, washed and infected as previously documented^71^. Cells were infected with test organisms and incubated for 4hr and 10hr. Lactate dehydrogenase release was determined using the Cytotox 96 cytotoxicity assay kit (Promega USA) as per manufacturers protocol.

### Catalase Activity Assay

Overnight cultures of bacterial strains were grown in LB broth, diluted 1:100 in the same medium and grown to an OD600 of 0.4. Catalase standards were prepared as per the manufacturers protocol using the OxiSelect™ Catalase Activity Assay Kit, Colorimetric (Cell Biolabs, inc). 20 μΙ of each serial dilution of overnight culture were added to 3 wells in a 96 well plate to allow for average readings for each sample. Plate absorbance was read at 520nm using FLUOstar Optima plate reader (BMG UK).

### Accessibility of biological resources

SCVs used in this study have been deposited at DSMZ under DSM 100776 - 100778.

## Acknowledgements

This work was funded by an MRC SCP3 fellowship grant G1000419 and by grant 8000-105-3 of the German Federal Ministry of Science and Education through the German Centre of Infection Research (DZIF) to J.O. We thank Simone Severitt and Nicole Mrotzek at the Leibniz Institute DSMZ, Germany, for excellent technical assistance.

## Author Contributions Statement

S.I, D.W, B.B and J.O conceived and designed the experiments and analysis. S.I, B.B, C.S performed the experiments. S.I, B.B, J.P.R.C and A.J.R analysed the data. T.J.E and H.K.B supplied novel reagents. D.W and S.I wrote the manuscript.

## Supplementary Figure

**Figure S1.**
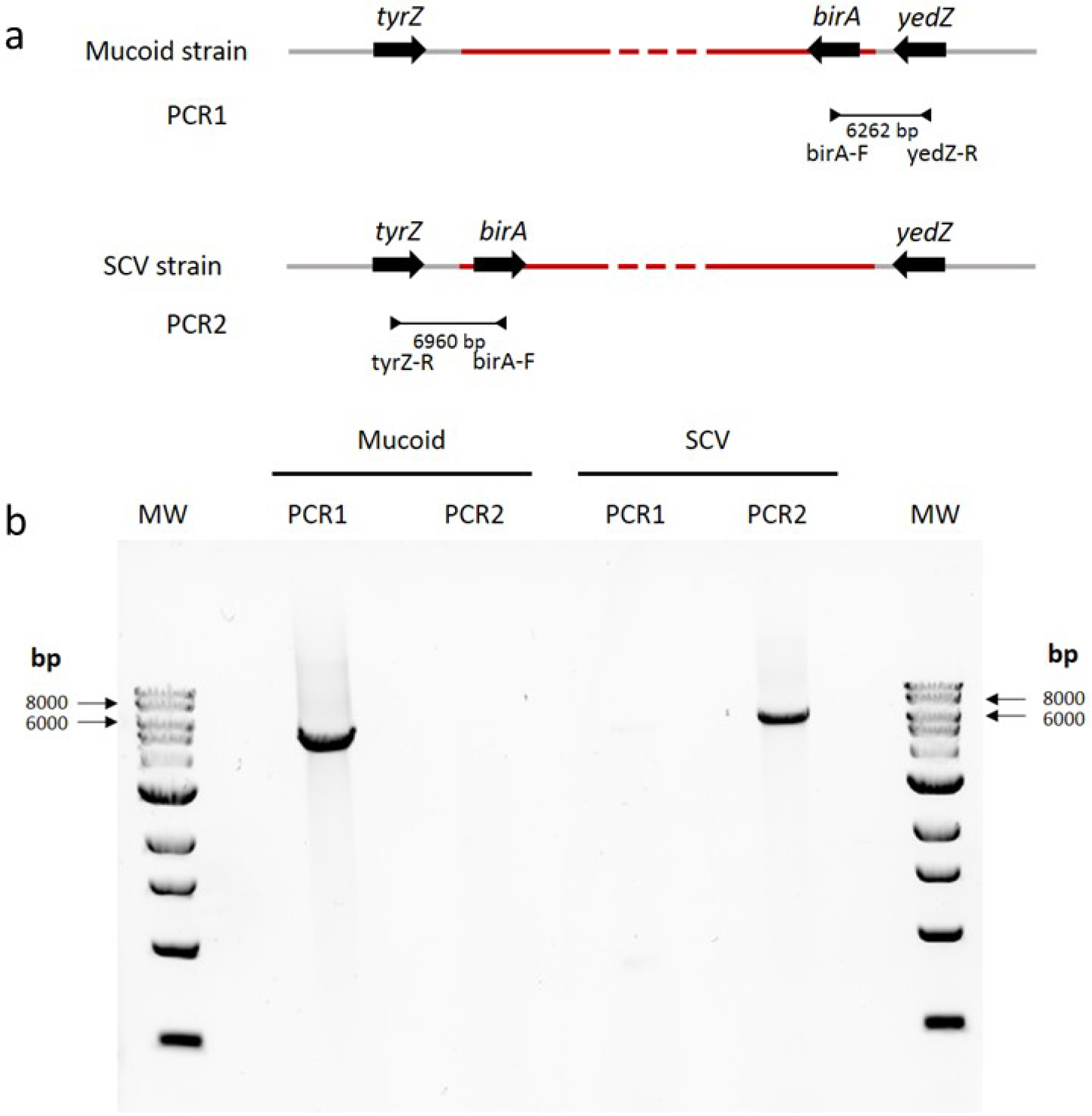
The mucoid strain, NHMucJan, isolated from the same chronic infection model as SCVJan does not contain a similar genome inversion. a) To determine if NHMucJan contains a similar genome inversion as SCVJan and SCVFeb we developed a PCR strategy using primers specific to *tyrZ* or *yedZ* which lie outside and adjacent to the inverted region in combination with primer specific to *birA*, which is adjacent to the rRNA genes within the inverted region. Only the primer pair expected to give a PCR product for each strain is shown. The inverted sequence in the SCV strain is highlighted in red. b) Consistent with the hypothesis that genome inversion drives conversion we observed a PCR product only with birA-F/yedZ-R primer pair for NHMucJan indicating that the genome structure in this region is identical to the parent strain NHMuc.

